# Combining niche modelling, land use-change, and genetic information to assess the conservation status of *Pouteria splendens* populations in Central Chile

**DOI:** 10.1101/026336

**Authors:** Narkis S. Morales, Ignacio C. Fernández, Basilio Carrasco, Cristina Orchard

## Abstract

**Background:** *Pouteria splendens* (lúcumo chileno) is an endemic shrub to the coastal areas of Central Chile classified as Endangered and Rare by the Chilean threatened species list, but as Lower Risk (LR) by IUCN. Based in historical records some authors have hypothesized that *P. splendens* originally formed a large metapopulation, but due to habitat loss and fragmentation these populations have been reduced to two main areas separated by 100 km, neither of both currently protected by the Chilean system of protected areas. Knowledge about this species is scarce and no studies have provided evidence to support the large metapopulation hypothesis. This gap of knowledge limits our availability to gauge the real urgency to conserve remaining *P. splendens* populations, which can generate tragic consequences in light of the increasing land-use change and climatic change that are facing these populations. In this study we combined niche modelling, land-use information, future climatic scenarios, and conservation genetics techniques, to test the hypothesis of a potential original large metapopulation, evaluate the role of land-use change in population decline, assess the threats this species may face in the future, and combine the generated information to re-assess its conservation status using the IUCN criteria.

**Results:** Our results show that locations with *P. splendens* are fewer than described in the literature. Results from the niche modelling and genetic analyses support the hypothesis of an originally large metapopulation that was recently reduced and fragmented by anthropogenic land-use change. Future climate change could increase the range of suitable habitats for *P. splendens* towards inland areas; however the high level of fragmentation of these new areas is expected to preclude colonization processes.

**Conclusions:** Based on our results we recommend urgent actions towards the conservation of this species, including (1) re-evaluating its current IUCN conservation status and reclassifying it as Endangered (EN), and (2) take immediate actions to develop strategies that effectively protect the remaining populations.

## Background

*Pouteria splendens* (A.DC.) Kuntze (lúcumo chileno, palo colorado) is an endemic shrub of the coastal areas of central Chile, and is the only representative of the Sapotaceae family in Chile (Muñoz & Serra 2006). *P. splendens* is an evergreen shrub, usually no taller than 2 meters, and with an intricate branching physiognomy. However, if conditions are favorable, individuals can develop longer trunks and reach more than 5 meters in height (Hechenleitner et al. 2005).

Historic records for *P. splendens* indicate a distribution that ranged from the coastal areas near the village of Huentelauquén (31°36’24’’S; 71°32’10’’O) in the north, to the beach resort of Algarrobo (33°22’50’’S; 71°40’57’’O) to the south, extending for near 200 km along the coastal range of central Chile. However, the current distribution is reported to be restricted to just two main populations isolated from each other by approximately 100 km (Hechenleitner et al. 2005). Among these, the northern area has the larger population, covering an important proportion of the coastal terrace between north of Pichidangui (32°08’19’’S; 71°31’46’’O) and south of Los Molles (32°14’15’’S; 71°30’52’’O). The southern area is represented by a few isolated groups of individuals located on coastal cliffs and ravines between the coastal villages of Laguna Verde (33°06”05’S; 71°40”00’O) and Quintay (33°11”38’S; 71°41”56’O).

Current knowledge about *P. splendens* populations is extremely scarce, and until now much of the literature is based on historical presence records and anecdotal reports. For example, although available literature suggests that current populations are the remnants of a larger metapopulation covering the entire range of the historic distribution (Hechenleitner et al. 2005; Muñoz & Serra 2006), at present there is no empirical evidence, besides the historical presence records, that supports this hypothesis. The lack of information regarding *P. splendens* is also illustrated by the inconsistencies among authors or institutions in relation to its conservation status. Whereas Squeo et al. (2001) and Hechenleitner et al. (2005) classified the species as critically endangered, the Chilean species inventory system has listed it as endangered (Ministerio de Medio Ambiente de Chile 2014), and the International Union for Conservation of Nature (IUCN) as lower risk/near threatened (International Union for Conservation of Nature 2014).

Among the main probable factors responsible for the reduction of *P. splendens* geographical range are the loss and fragmentation of original habitat due to the development of coastal resorts, towns and cities that now cover most of the original range of its distribution (Hechenleitner et al. 2005). Alongside habitat loss, the local extinction of large animals capable to disperse *P. splendens* seeds may have reduced the probability of these seeds to find new suitable sites for germination (Henríquez et al. 2012; Sotes et al. 2013). Additionally, the increasing frequency of droughts experienced by these areas due to climate change (Quintana 2012; Schulz et al. 2012) may have reduced *P. splendens* germination rates (Hechenleitner et al. 2005; Sotes et al. 2013) and increased its vulnerability to local extinctions due to an increment in fire regimes (Montenegro et al. 2004). Climatic conditions are expected to keep changing in all the range of *P. splendens* distribution (CONAMA 2006; Nuñez et al. 2009), which may generate additional burdens for the preservation of remaining populations. Moreover, genetic diversity could decrease in fragmented populations as a consequence of genetic drift, inbreeding, bottlenecks, and founder effects, which together with demographic and environmental fluctuations may determine the persistence of populations in the long run (Lammi et al. 1999).

Even though remaining populations are increasingly threatened by human actions, at present none of the *P. splendens* populations have been covered by the Chilean system of national parks and protected areas (SNASPE), nor have they been included by any kind of public or private effective protection or conservation strategies. This lack of action has occurred even though the northern range of *P. splendens* distribution was officially recognized as an urgent site for conservation one decade ago (CONAMA-PNUD 2005).

The few studies available for *P. splendens* have been focused in understanding the species autoecology (i.e., Sotes et al. 2006; Nuñez et al. 2009; Henríquez et al. 2012; Sotes et al. 2013), and until now there have been no studies at larger scales that may provide answers to the original metapopulation hypothesis. This lack of information is not only limiting our understanding about potential causes of the current fragmented distribution of *P. splendens* populations, but also our ability to gauge the urgency to develop conservation efforts.

The combination of spatially explicit ecological tools, such as species niche modelling (e.g., Phillips et al. 2006; Phillips & Dudík 2008; Mateo et al. 2011) and conservation genetics techniques (e.g., Frankham 2005; Freeland 2005) may provide critical information regarding species original distribution. Furthermore, the increasing availability of spatial land-use databases can offer key insights into the main past causes of population degradation, whereas the development of future climate models may provide valuable information regarding population potential future trends. Combining and evaluating this information should be taken as major task prior to developing conservation strategies for threatened species (Opdam & Wascher 2004).

In order to reduce the knowledge gap regarding *P. splendens* past and current populations, and to provide insightful knowledge that can be used for developing planning efforts towards its conservation, in this study we had five main objectives: (1) estimate the potential original distribution range of *P. splendens* by using a niche modelling approach, (2) evaluate potential trends of *P. splendens* populations under a probable future climate change scenario, (3) evaluate the role of land-use change in current and future species distribution, (4) assess intra and inter population genetic diversity in order to evaluate the hypothesis of an original large continuous population, and (5) assess its current IUCN conservation status using this generated data.

## Methodology

### Study Area

The study area covered the coastal range of North Central Chile (Fig. 1), which is the historical reported distribution range of *P. splendens* (Hechenleitner et al. 2005; Sotes et al. 2013). This area can be broadly characterized as having a Mediterranean climate, with colder temperatures and rainy periods during winter months, and warmer and drier conditions during summer period (Luebert & Pliscoff 2006). Annual average precipitation within this area ranges from ∼180 mm to ∼400 mm in an increasing gradient north-to-south, but monthly distribution of precipitation follow the same pattern in the whole range, with July being the wettest month and February the driest. Temperatures are relatively similar for the entire study area, with a mean annual temperature of ∼13°C, where July is the coldest month (∼10.5°C) and January the warmest (∼17°C) (Dirección Meteorológica de Chile 2001). Vegetation within this area is characterized by a predominance of shrub species that are directly influenced by oceanic conditions, with only occasional presence of forest patches, mainly in creeks and valleys facing southern slopes (Gajardo 1994).

**Figure 1.**
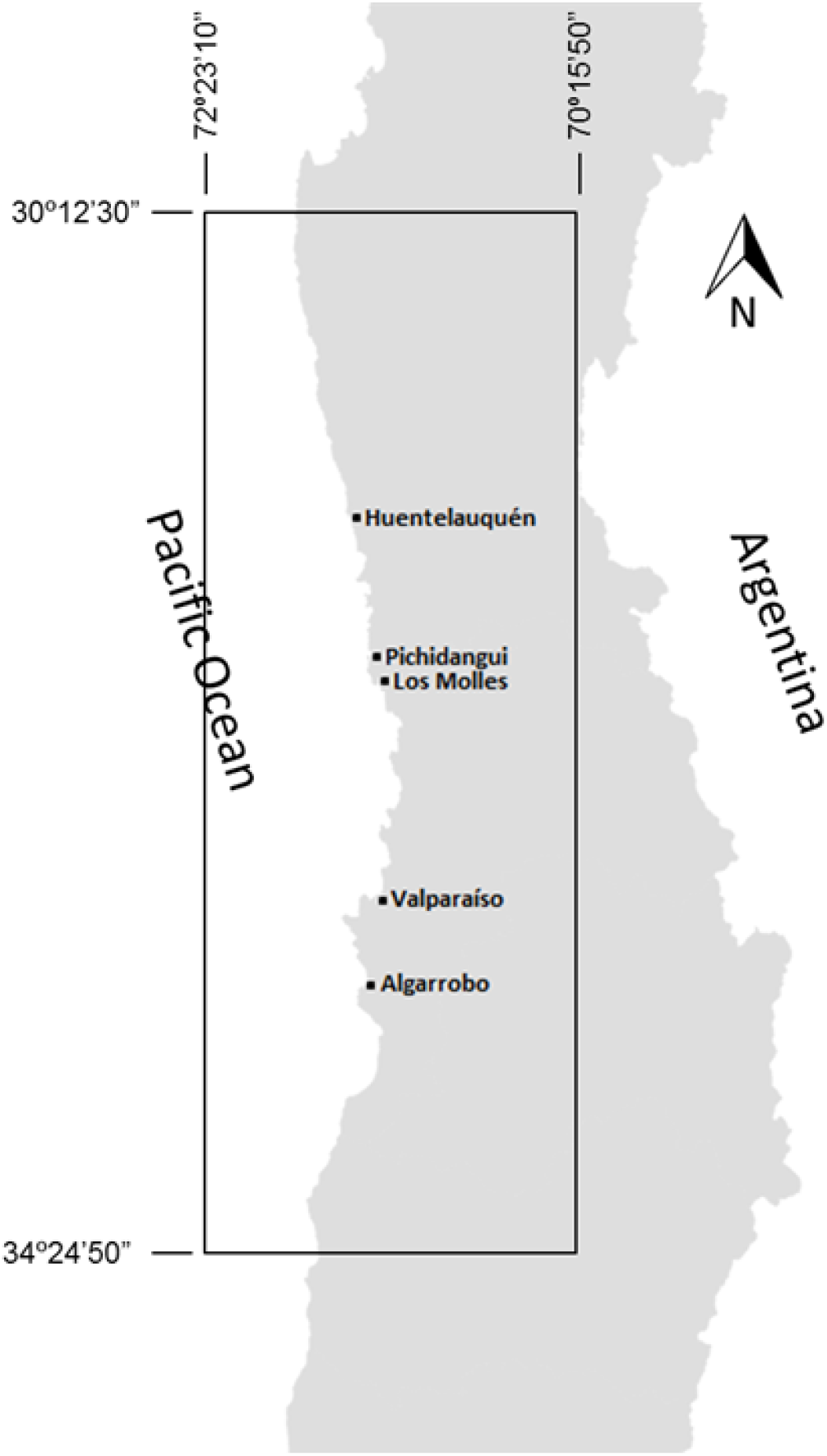
Area of Study. The area of study covers the coastal areas of central Chile. Historical presence records for *P. splendens* goes from the coastal village of Huentulauquén to the beach resort of Algarrobo. Dots respresent recorded presence for the species, filled dots are those for which we found in-site current presence of individuals. The black rectangle indicates the area magnified in the binary distibution maps shown in Fig. 2.

### P. splendens population surveys

A total of 22 presence data points of *P. splendens* were used in our study. From these, 21 points were collected from herbarium records of the Chilean National Herbarium (SGO) and from the University of Concepción Herbarium (CONC). Records were georeferenced and visited on several field trips during the spring of 2010. We also visited other potential sites having the species, basing our search on satellite imaging interpretation and personal communications. A 22^nd^ presence point was registered during the field trip, representing a small population not previously recorded (see supplementary data). In all the sites we found established individuals we did a rough estimation of abundance by a visual prospection in a ∼30 minutes walking distance.

### P. splendens potential habitat modelling

To model the potential habitat distribution of *P. splendens* we used the principle of maximum entropy, which is part of the software MaxEnt (Phillips et al. 2006). MaxEnt estimates the probability of occurrence of the species of interest using presence only data and a group of environmental variables. We decided to use MaxEnt because it has been proven to be one of the best performing modelling methods available (Elith et al. 2006), it is reliable even with a small number of samples (Pearson et al. 2007), and is freely available. All the habitat distribution models were performed using MaxEnt version 3.3.3k (http://www.cs.princeton.edu/∼schapire/maxent).

Environmental variables for modelling were gathered from the WorldClim database. This database consists of a set of 19 global climatic layers generated by the interpolation of global climatic data such as average and seasonal temperature and precipitation (Hijmans et al. 2005). We used the climatic data corresponding to the period from 1950 to 2000, with a resolution of approximately 1 Km^2^ (30 arc-sec).

Considering the level of spatial resolution of the environment data layers, and to avoid pseudo replication, we checked that our 22 *P. splendens* presence records were at least 1 km^2^ apart. Additionally, to reduce the potential of model over-parameterization (Merow et al. 2013), we performed (Pearson) correlation analysis of the 19 bioclimatic variables, evaluating all pairs of variables with levels of correlation higher than 0.8 (r >0.8), and keeping only those variables that were biologically significant or easier to interpret. The correlation analyses were performed using the software ENMTools version 1.4.4 (Warren et al. 2010, Warren & Seifert 2011). After the correlation analysis only 8 out the 19 bioclimatic variables were used for distribution modelling (Table 1).

**Table 1.** Bioclimatic variables from WorldClim database. The variables in bold type were chosen to model the distribution of *P. splendens* after Pearson correlation analysis. The most relevant variables explaining the current distribution of *P. splendens* are shown in the right column.

| Variable | Model contribution (%) |
| --- | --- |
| Annual Mean Temperature (Bio 1) | - |
| Mean Diurnal Range (Bio 2) | - |
| **Isothermality (Bio 3)** | **22.4** |
| **Temperature Seasonality (Bio 4)** | **14.4** |
| Max Temperature of Warmest Month (Bio 5) | - |
| **Minimum Temperature of Coldest Month (Bio 6)** | **62.7** |
| **Temperature Annual Range (Bio 7)** | **0.3** |
| Mean Temperature of Wettest Quarter (Bio 8) | - |
| Mean Temperature of Driest Quarter (Bio 9) | - |
| Mean Temperature of Warmest Quarter (Bio 10) | - |
| Mean Temperature of Coldest Quarter (Bio 11) | - |
| **Annual Precipitation (Bio 12)** | **<0.1** |
| Precipitation of Wettest Month (Bio 13) | - |
| **Precipitation of Driest Month (Bio 14)** | **0.1** |
| Precipitation Seasonality (Bio 15) | - |
| Precipitation of Wettest Quarter (Bio 16) | - |
| Precipitation of Driest Quarter (Bio 17) | - |
| **Precipitation of Warmest Quarter (Bio 18)** | **<0.1** |
| **Precipitation of Coldest Quarter (Bio 19)** | **<0.1** |

We did not use the default configuration provided by MaxEnt for small samples (i.e., auto features) as recent studies have shown that this configuration would not be the most appropriate in some cases (e.g., Merow et al. 2013; Syfert et al. 2013), especially when dealing with a small number of samples (< 20 - 25) (Shcheglovitova & Anderson 2013), as it was in our case. Therefore, to determine the optimal parameters to configure MaxEnt we compared different models with a combination of the “feature class” and “regularization multiplier” parameters recommended when dealing with small number of samples (Shcheglovitova & Anderson 2013). MaxEnt provides different types of restrictions (“feature class”) in the modelling stage such as lineal (L), quadratic (Q), product (P), threshold (T) and hinge (H) (Phillips et al. 2006). We used the following combinations of these features: L, H, LQ, LQH. The regularization multiplier values used were based on Warren and Seifert (2011): 1, 2, 5, 10, 15 y 20. All the models were compared using the corrected Akaike information criterion (AICc) (Warren & Seifert 2011), using for this process the software ENMTOOLS version 1.4.4 (Warren et al. 2010). The results suggested that the best combination of parameters were LQ with a regularization multiplier of 1 (see supplementary data).

Once the optimal parameters were determined, we analyzed the performance of the model of choice using the method proposed by Pearson et al. (2007) for models with small a number of samples. The method consists on a jack-knife procedure that removes one presence point from the original dataset (n=22) to subsequently run the model with the remainder (n – 1, 21 points) presence points. This process is repeated with all the available presence data points, generating 22 different models. Later, the model performance was analyzed using a prediction success rate and a *p* value using the software “pValuecompute v.1.0” provided by Pearson et al. (2007).

We produced binary maps using two different thresholds to define the suitable vs. non-suitable habitat. We used the “maximum sensitivity plus specificity logistic (MSS)” and the “10 percentile training presence logistic (10PL)” thresholds (values of 0.203 and 0.298 respectively). We used these parameters as they are among the two most commonly used thresholds for creating binary suitability maps for species distribution with MaxEnt (e.g., Liu et al. 2005; Pearson et al. 2007; Contreras-Medina et al. 2010; Vale et al. 2014).

### P. splendens potential habitat modelling under climate change

To predict potential changes in the habitat distribution of *P. splendens* under a hypothetical climate change scenario, we re-projected the model generated in the previous methodological section using a new environmental layer from the period (2041-2060). We used the climatic projections from the General Circulation Model (GMC) from the United Kingdom Meteorological Office Hadley Centre known as HadGEM2-ES (Jones et al. 2011). The new climatic layers were downscaled and calibrated (bias corrected) using WorldClim 1.4 as baseline “current” climate (Hijmans et al. 2005). We used an optimistic approach by using the RCP 2.6 scenario of representative concentrations pathways (RCP) of greenhouse gases (Moss et al. 2010). The RCP 2.6 scenario assumes that the global greenhouse emissions will have their maximum concentration between the years (2010 - 2020), declining substantially after this period. We used the same bioclimatic variables utilized to build the habitat distribution model under the “current” climatic conditions. Binary maps for projected future potential distribution were generated using the same thresholds and approaches used to build the current conditions maps.

### Land Use-Change mapping

To estimate the historical land-use change that has occurred within our study area we used the Chilean Forest Inventory (freely available at http://sit.conaf.cl), which is the most comprehensive and recently updated geodatabase of land-use types of Chile (CONAF 2011). From this vector-based geodatabase we identified and grouped all the land-use categories that were explicitly indicating a replacement of original natural habitat, including urban areas, industrial areas, mining areas, agriculture lands, and silvicultural lands. All these polygons were grouped under a single category (i.e., land-change) and rasterized in a 100 m pixel size grid to create a land-change layer compatible with the Maxent lattice output. This generated layer was then subtracted from the niche modelling layers to map and calculate the extension of habitat loss. Because we did not have an accurate way to estimate the future trends in land-use change, we decide to use a heavily conservative approach to map future potential distribution under land-use change by assuming that current distribution and extension of anthropogenic land-use will remain constant.

### Genetic diversity of current populations

We used Inter Simple Sequence Repeat (ISSR, Gupta et al. 1994; Zietkiewicz et al.1994) to evaluate the level of inter and intra genetic diversity of current *P. splendens* populations. We collected samples of young leaves from a total of 121 individuals from three different areas (Table 2). These areas represent the northern, southern and largest populations for which we found individuals, and were analyzed as three different populations for the genetic analysis. All sampled individuals were georeferenced and their leaves frozen in liquid nitrogen and transported to perform DNA extraction and amplification (through PCR) according to Carrasco et al. (2007).

**Table 2.** Genetic variability parameters obtained from 147 bands in 121 individuals of *P. splendens.* N: number of individuals sampled per area; I: Shannon’s index; He: expected heterozygosity; P (%): percentage of polymorphic loci.

| Area | N | I | He | P (%) |
| --- | --- | --- | --- | --- |
| Palo Colorado (PC) | 48 | 0.40± 0.01 | 0.24± 0.01 | 100 |
| Pichidangui-Los Molles (PM) | 43 | 0.35± 0.01 | 0.21 ±0.01 | 99.32 |
| Caleta Quintay (CQ) | 30 | 0.35± 0.01 | 0.21 ±0.01 | 98.64 |

We tested a set of 16 primers (set ISSR 100/8, Biotechnology Laboratory from University of British Columbia, Vancouver), from which five primers were selected to perform the PCR. The basic characteristics of the primers that gave positive results are shown in supplementary material. Amplified genetic material was run by electrophoresis (2% agarose gel) and patterns were visualized and manually recorded (i.e., band presence or absence) for each individual.

We calculated the percentage of polymorphic ISSR loci (P%) and expected heterozygosity (He) (Nei 1973) by using GenAlex 6.4 computer software (http://biology-assets.anu.edu.au/GenAlEx/Welcome.html; Peakall & Smouse 2012). The relative degree of phenotypic diversity was measured using the Shannon’s index.

We estimated the genetic differentiation between populations by using AMOVA Φst (Excoffier et al 1992). We also performed a Principal Coordinate Analysis (PCoA) to evaluate the patterns of clustering based on the binary genetic distance displayed by the GenAlex Software (Peakall & Smouse 2012). In order to analyze the genetic relationship among sampled individuals, we used the Structure Software Version 2.1 (http://pritchardlab.stanford.edu/structure.html), which is a method that assigns individuals from the entire sample to *k* clusters in a way that the Hardy Weinberg equilibrium are maximally explained (Pritchard et al. 2000). We used Evanno et al. (2005) methodology to estimate the *k* number of clusters by the Structure Software, using a threshold of 0.8 as the membership probability value (*Q*) to assign individuals to a specific cluster.

### Assessment of the conservation status of P. splendens according to IUCN criteria

We used the IUCN Red List Categories and Criteria version 3.1 that includes five criteria to assess the species level of threat (International Union for Conservation of Nature 2001). The criteria are: (a) reduction in population size; (b) small geographic range; (c) small population size and decline; (d) very small or restricted population; and, (e) quantitative analysis of extinction risk. We combined existing information with the data generated in our study to re-evaluate the conservation status of *P. splendens*.

## Results

### P. splendens population surveys

After the field campaigns we were able to confirm the current presence of *P. splendens* in only nine sites, which were aligned to the species distribution range reported in the literature. From these, eight corresponded to sites previously reported to contain the species, and one to a site not recorded before. The northern and southern sites corresponded to relatively small and isolated populations (<100 individuals), whereas the sites reported between Pichidangui and Los Molles corresponded to an apparently relatively large (thousands of individuals) and continuous population (see supplementary material for sites information).

### P. splendens potential habitat modelling

The most relevant bioclimatic variables that explained *P. splendens* habitat distribution were “Minimum Temperature of Coldest Month” and “Isothermality”, accounting for 62.7% and 22.4% of the predicted distribution respectively (Table 1). A performance test for the predicted distribution of *P. splendens* showed that the model had a good predicting performance, with a success of 86% (*p* < 0.001) for the MSS approach, and 76% (*p* < 0.001) for the 10PL approach. The area for *P. splendens* predicted by the model using both of these techniques was mostly restricted to coastal plains, ravines and valleys within its historical range. Even though there was a 33.26% difference in the total predicted suitable habitat (Table 3), both techniques identified three main areas that concentrated the large part of suitable habitat. The largest of these areas was located in the central part of the predicted distribution, an intermediate area was located in the southern range, and the smallest was in the northern section of the distribution (Fig. 2).

**Table 3.** Potential habitat, land-use change, and habitat loss of *P. splendens* under current climatic conditions (year 1950 to 2000) and future climate change scenario (year 2041 to 2060) with different thresholds; maximum sensitivity plus specificity (MSS) and 10 percentile training presence (10PL).

| Threshold | Current |  |  | Future |  |  |
| --- | --- | --- | --- | --- | --- | --- |
|  | Original<br>(ha) | Land-use<br>change (ha) | Habitat<br>lost (%) | Original<br>(ha) | Land-use<br>change (ha) | Habitat<br>lost (%) |
| MSS | 140,267 | 99,003 | 29.42 | 614,529 | 420,280 | 31.61 |
| 10PL | 93,617 | 65,238 | 30.31 | 521,533 | 362,364 | 30.52 |
| Difference<br>(%) | 33.26 | 34.10 |  | 15.13 | 13.78 |  |

**Figure 2.**
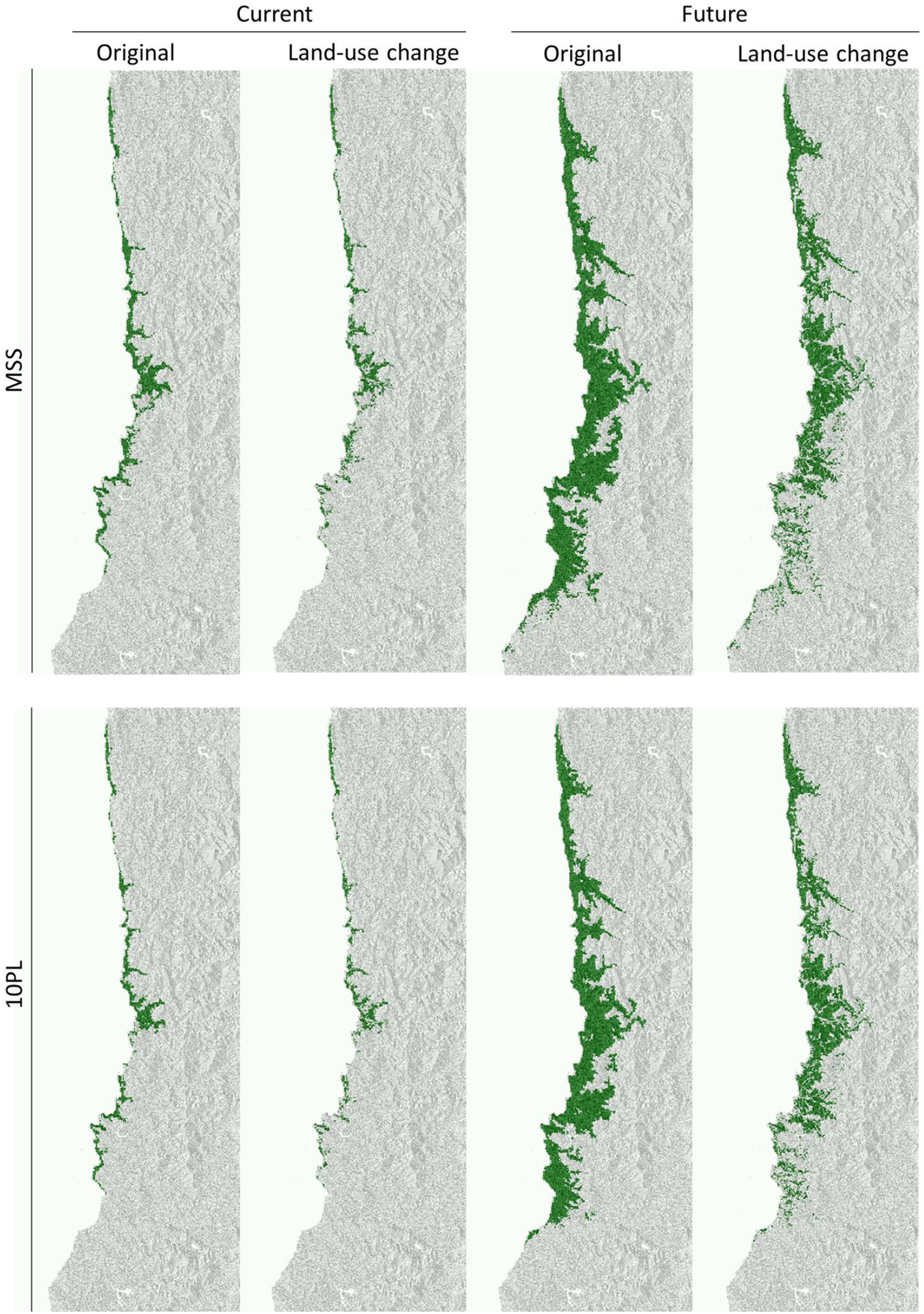
Modelled distribution maps. Maps show potential habitat (green) of *P. splendens* with or without land-use change under current climate (year 1950 to 2000) and future climate change scenario (year 2041 to 2060). Two different thresholds were used two generate the binary maps; maximum sensitivity plus specificity (MSS) and 10 percentile training presence (10PL).

### P. splendens potential habitat under climate change

Results from the predicted distribution under climate change scenario show that the latitudinal (north-south) range for *P. splendens* is not expected to change much in relation to the predicted distribution under current conditions (Fig 2). However, there is a large effect on the longitudinal extension of populations, with future populations not only located in the coastal range, but also spreading further east through the valleys towards the Chilean central valleys. This increase in distribution also generates an important reduction in the differences between the total potential areas predicted by the two aforementioned techniques when compared to the predicted total area under current climatic conditions (Table 3).

### Estimation of land-use change effects on P. splendens potential distribution

When areas subjected to land-use change are subtracted from maps of potential distribution a clear sign of habitat loss and fragmentation appears (Fig. 2). The largest extent of this habitat loss was located in the coastal range of *P. splendens*’ potential distribution and was more noticeable in the central and southern range of its distribution. The effect of habitat loss on potential distribution under current climatic conditions was larger in absolute values (ha) in the 10PL approach in comparison to the MSS (Table 3). However, in relative terms the percentage of habitat loss was practically the same, with nearly 30% of the potential area of distribution being lost by anthropogenic land-use change.

In the case of projected distribution under future climatic conditions there was a considerable increase of land-use change effects in absolute values in comparison to the same factor under current conditions. However, surprisingly when the effects of land use change were analyzed in relative terms, proportion lost values were not different for current and future scenarios, and were slightly higher than 30% for both future threshold approaches (Table 3).

### Genetic diversity of P. splendens

We obtained a total of 157 polymorphic loci with fragments ranging between 260 and 2500 base pairs depending on the used primer. The level of genetic diversity was similar between individuals from the three sampled areas (hereafter populations). The level of polymorphism varied between 98 to 100% for the studied loci. Heterozygosity and Shannon’s diversity index ranged between 0.21-0.24 and 0.35-0.40 respectively.

Results from the AMOVA analysis indicated that 97% of the genetic diversity was distributed within populations, and only a small proportion of genetic diversity could be assigned to differences between populations (ϕst= 3%, *p*=0.01). In addition, the Principal Component Analysis (PCA) did not show any grouping patterns that may reveal presence of genetic differences between sampled populations (see supplementary data).

The Bayesian Clustering analysis performed through the Structure Software using both Evanno et al. (2005) and Pritchard et al. (2000) criterion resulted in a *k* number of 3 clusters (Figure 3). The resulting clusters did not show a genetic structure aligned with the spatial distribution of sampled populations. The first cluster included 23 individuals (15 from “Palo Colorado (PC)”, 6 from “Pichidangui-Los Molles (PM)”, and 2 from the “Caleta Quintay (CQ)” population). The second cluster was comprised of 19 individuals (4 from PC, 7 from PM, and 8 from the CQ population). The third cluster included only nine individuals, all of them from the PM population. The remaining 70 individuals (58% of sampled individuals) showed *Q* values lower than 0.8, and therefore considered as a genetically mixed group.

**Figure 3.**
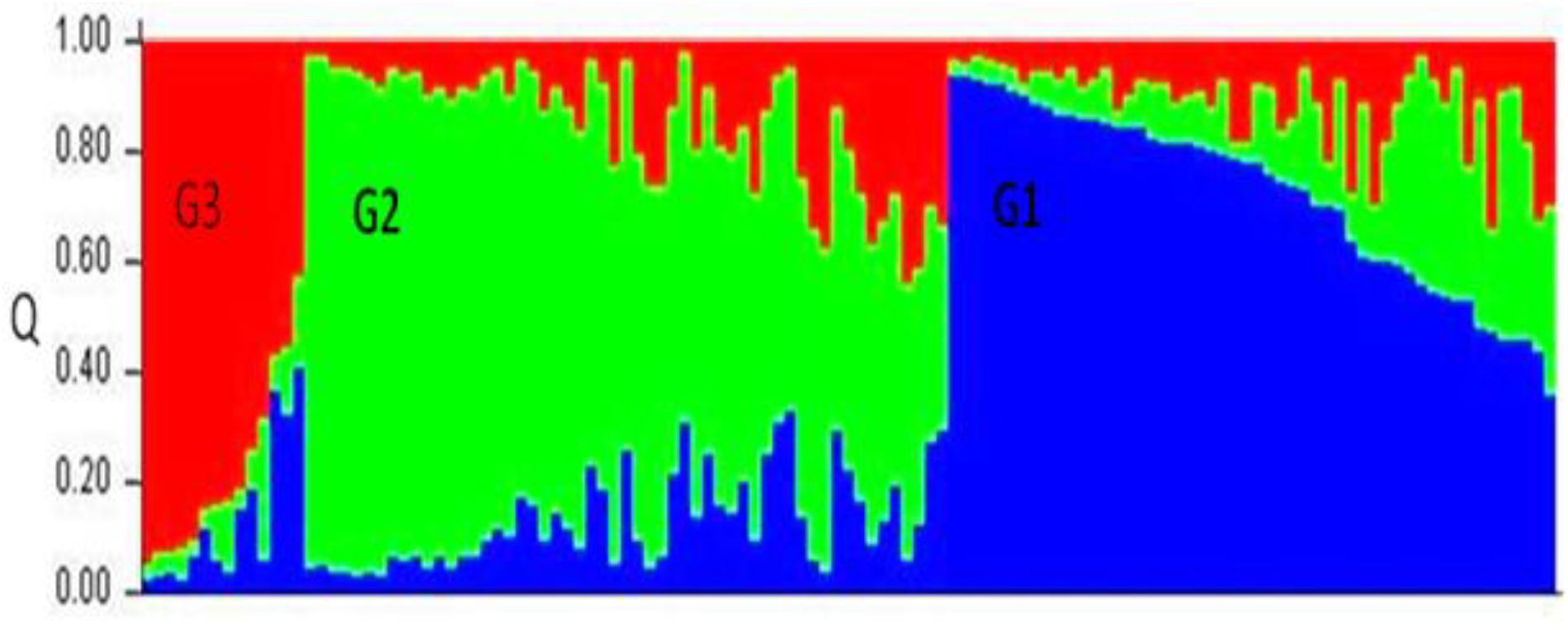
Bar plot of *Q* values. Genetic diversity analysis by using the structure software shows three clusters. Blue cluster (G1) included 23 individuals; green cluster (G2) included 19 individuals, and red cluster (G3) contained 9 individuals. 70 individuals of *P. splendens* did not pertain to any of the three clusters (*Q* values <0.8).

### A*ssessment of the conservation status of P. splendens according to IUCN criteria*

Results from our study and available information in the literature allow us to evaluate this species for only three out of the five criteria listed by the IUCN (see method section). We could not evaluate criteria (a) because the lack of data about regeneration length. We could not evaluate criteria (e) because we did not have enough data to perform extinction risk analysis such as PVA. Therefore we only assessed *P. splendens* conservation status by evaluating criteria (b), (c), and (d) (Table 4). To perform the analysis we used the estimated habitat loss presented in Table 3 and the rough estimation of the number of individuals present in the validated sites (*n* = 1650, see supplementary material). We complemented this data with our genetic diversity analyses that suggested that the populations sampled for the analysis were part of one original metapopulation now fragmented. Also, autoecology of the species was considered in the assessment of the level of threat, such as lack of regeneration and seed dispersion constraints (i.e., lack of seed dispersers). In addition, we incorporated references that mentioned threats to *P. splendens* and its habitat (i.e. fires). Moreover, we estimated habitat loss in the next decades in the area of occurrence of *P. splendens* because the probable development of an urbanization project that potentially will destroy 1700 ha of this species habitat. The results of the threat assessment are summarized in Table 4. Based on our results we recommend classifying *P. splendens* as an Endangered (EN) species.

**Table 4.**
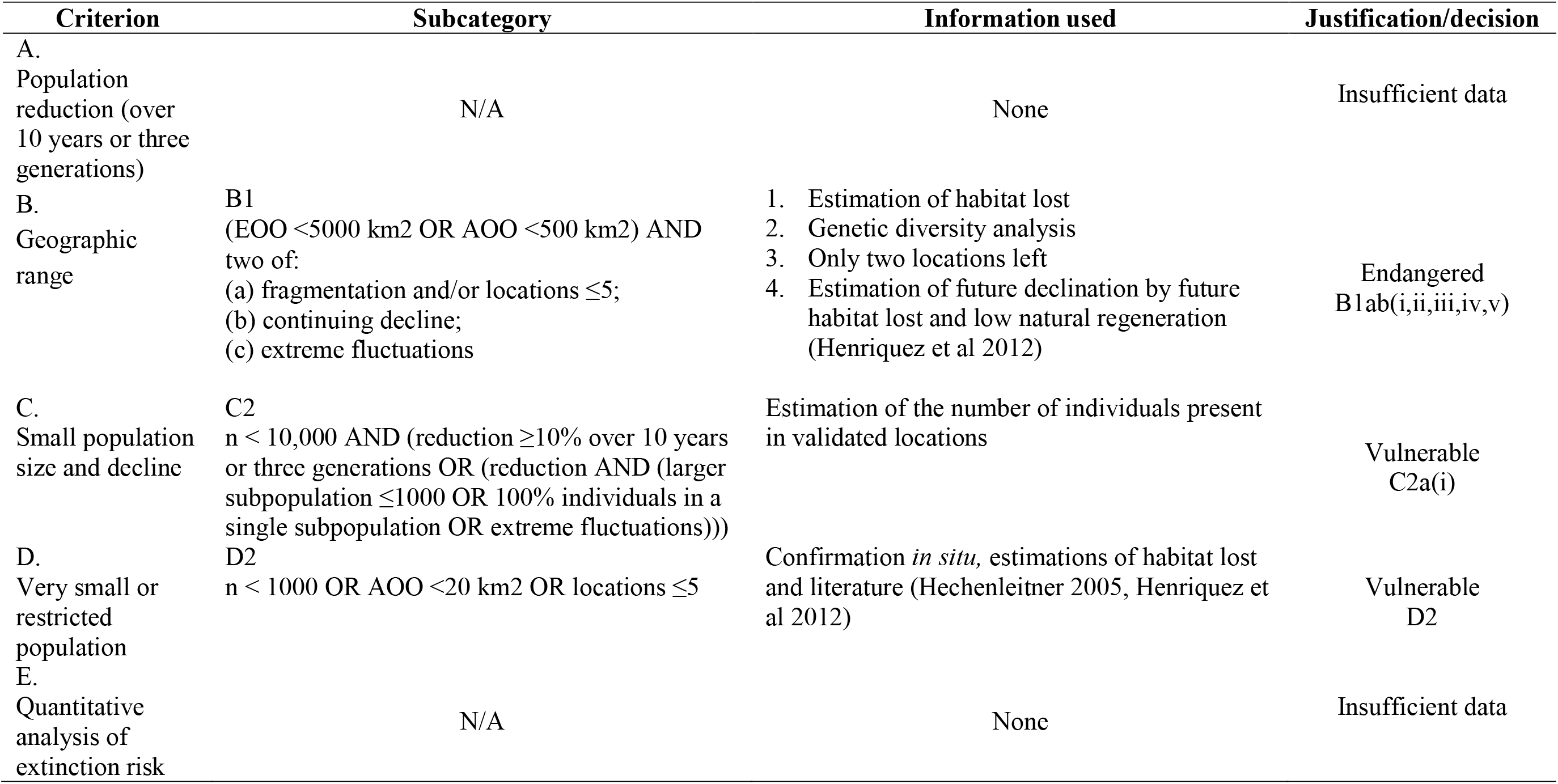
IUCN Red List criteria that apply to *P. splendens* with description of the information used in the assessment and its applicability in the present study. Abbreviations within the table; EOO: Extent of Occurrence, AOO: Area of Occupancy, n: mature individuals (Based on Cardoso et al. 2011).

## Discussion

This is the first attempt to study the distribution of *P. splendens* throughout its current and historical range. Furthermore, this study could be considered as one of the first efforts to generate landscape-level knowledge of this species, and the first aiming to provide evidence to evaluate the hypothesis of an original large and continuous population that has been severely reduced and fragmented by historical land-use change. In addition, it is probably the first effort to assess the conservation status of this species by relying on quantitative data.

Increase the knowledge about understudied taxa with potential conservation problems is a key task for evaluating the real urgency to take conservation actions in the short-term (McKinney 1999). However, this implies several challenges due to the scarcity of available and/or reliable background data. One of the primary challenges we faced in developing this study was the difficulty in finding individuals at locations where historical records show its presence. From a total of 22 historical recorded presence points, we could only validate the presence of *P. splendens* in eight sites. Some of the historic records are very old (e.g., La Palma 1926), so we expected that some populations could have gone extinct or have declined in density and extent. Therefore, it is probable that present densities were strongly reduced in some sites, which could preclude us to detect the individuals that were reported in the records. Furthermore, because the nature of the available records (i.e., historical herbarium lists), some locations were not properly georeferenced and were described simply as a locality or geographic landmark (e.g., “Chivato’s ravine”). This probably decreased our success rates of finding individuals representing the populations listed in the historical records. However, we also visited several points not described in the literature that we thought would be suitable for the presence of *P. splendens*, with no individuals found except from one site. The difficulty of finding populations of this species within their historical range of distribution support the urgent call for conserving remaining populations made by Hechenleitner et al. (2005).

### Current and future suitable habitats

The small number of presence records for *P. splendens* was another challenge we had to deal in our study. Although Maxent has been reported to perform well even with small number of samples (Pearson, 2007), we took additional actions by tuning the model default setting to optimal levels as this procedure has shown to considerably increase the model performance when dealing with small numbers of presence data (Shcheglovitova & Anderson 2013).

The modelled potential original habitat distribution of *P. splendens* is concentrated in coastal areas between the village of Huentelauquén and the beach resort of Algarrobo, which is in accordance with the historical distribution described by other authors (e.g., Hechenleitner et al. 2005; Muñoz & Serra 2006; Sotes et al. 2013). However, the model also generated a small isolated strip of suitable habitat north of Huentelauquén, for which we do not have registers of known populations. Absence of registries for *P. splendens* in this northern area could be just due to a lack of information because it corresponds to a coastal plain far from towns and roads, and of difficult access. However, the area is also coincident with the presence of the Fray Jorge National Park (30°39’50’’S; 71°40’50’’O), for which there are detailed flora inventories (see Squeo et al. 2004), but no registries of *P. splendens*. Therefore, our result from the niche modelling needs to be carefully interpreted, because the lack of other potential important variables in our model (e.g., topography, soil, dispersion barriers) may result in an overestimation of suitable habitats.

Results from MaxEnt modelling using the climatic change scenario showed that *P. splendens* would spread towards the inland areas of Central Chile, considerably expanding its distribution range in comparison with its distribution under current climate conditions. This trend could be explained because the increase of temperatures between 2-3°C that is projected to occur in Central Chile due to climate change (CONAMA 2006; Nuñez et al. 2009). The *Pouteria* genus have a tropical original (Silva et al. 2009) and *P. splendens* is considered a relict species from an ancient warmer climate (Gajardo 1994; Francois 2004). Therefore, as temperature of the coldest month is the most important variable for this species, climate change may increase this temperature in inland territories, increasing potential suitable areas for this species in the future.

However, the projected potential distribution under climate change may overestimate the suitable areas for *P. splendens* because our model did not account for the effect of the marine fog commonly present in these coastal areas. Marine fog may have an important influence in the viability of *P. splendens* populations since it provides a relatively constant source of water (Hechenleitner et al. 2005; Muñoz & Serra 2006; Henríquez et al. 2012). Furthermore, projected climate change it supposed to reduce precipitation in inland areas of Central Chile (CONAMA 2006; Nuñez et al. 2009), which may reduce the germination rates of this species (Sotes et al. 2013). Therefore, we expect that the suitable areas will increase under a climate change scenario, but probably in a magnitude much smaller that is shown by the model.

In terms of the effects of land-use change in *P. splendens* distribution, our results show that the current potential habitat distribution is at least 30% smaller than the original potential distribution. This was regardless of the threshold used to estimate the species distribution. This percentage of potential habitat loss appears not to be so substantial if we take into account that this land-use change started more than 200 years ago, and that today nearly one million people live within this area. However, maps show that land-use change has also resulted in high levels of habitat fragmentation, which together with habitat loss, are considered as the main causes of biodiversity loss in Mediterranean habitats (Soulé et al. 1992; Vogiatzakis et al. 2006; Cooper et al. 2012).

Fragmentation of the original habitats imposes new conditions to the remaining communities, increasing edge effect and reducing the core area suitable for species (Fahrig 2003; Ewers & Didham 2006; Ewers et al. 2007). Furthermore, in central Chile fragmentation can increase the frequency of disturbances by fires (Montenegro et al. 2004; Fernández et al. 2010). Therefore, even though the modelled potential distribution suggests that nearly 70% of the areas suitable for *P. splendens* still are considered as remaining natural habitats, the additional effect of fragmentation may have caused an extreme reduction of the original populations. These current levels of habitat fragmentation also suggest that even assuming that under the future climate scenario suitable habitats for *P. splendens* will increase; barriers to dispersion may restrict the colonization processes (Engler & Guisan 2009; Henríquez et al. 2012).

### Genetic diversity

The analysis of inter simple sequence repeat (ISSR) diversity revealed that remaining *P. splendens* populations still maintain a high level of intra-population genetic diversity. The genetic variability values for *P. splendens* were similar to those reported by Nybom (2004) for plants species using dominant molecular markers, and are consistent with values observed for other perennial-endemic-alogamous-animal-dispersed species (Nybom 2004). In relation to the inter-population genetic diversity, the low levels of genetic differentiation between populations detected by AMOVA (ϕ_ST_ = 0.03) are notably lower than those observed for other plants species using dominant markers (Nybom, 2004), and also for populations of other natives Chilean tree species such as *Araucaria araucana* (ϕ_ST_ = 0.13; Bekessy et al. 2002) and *Pilgerodendron uviferum* (ϕ_ST_ = 0.19; Allnutt et al. 2003). The large intra and small inter-population genetic variability was also supported by the results from the Bayesian model-based clustering analysis. The analysis suggests the presence of three main clusters; however only 42% of the total analyzed individuals were assigned to a cluster, whereas the remaining 58% did not present enough genetic differences to be considered as part of any of the three clusters, and therefore were considered a mixed group.

Naturally rare or narrowly distributed plant species generally maintain lower within and higher between population genetic diversity than widespread species (Gitzendanner & Solis 2000; Aguilar 2008); however this was not the case for *P. splendens*. Among the main factors explaining a high genetic diversity in rare plant species are the recent reduction and isolation of populations, or the presence of specific mechanisms that promote gene flow between populations, such as animal or wind mediated outcrossing pollination and seed dispersal (Luan et al. 2006; Millar et al., 2014). In the case of *P. splendens* there is no available information about the pollination mechanism, but observation from other *Pouteria* species report self-pollination and insect-mediated pollination (Jordan 1996). None of these mechanisms could explain the genetic diversity patterns found within and between *P. splendens* populations. In relation to seed dispersal, the fruit size (2-3 cm in diameter, Hechenleitner et al. 2005) suggests that seed dispersal of *P. splendens* is probably mediated by large animals (Sotes et al. 2013). However, the long extirpation of large animals from the actual range of *P. splendens* distribution make improbable that this mechanism could be related to the genetic diversity of current populations.

Therefore, the high level of genetic variability within populations, the admixture patterns, and the weak genetic structure between populations, suggest that current *P. splendens* populations might be the result of a recent process of loss and habitat fragmentation. As time since fragmentation is a key factor related to the loss of genetic diversity in isolated populations (Aguilar et al. 2008), we may expect to see a decrease in intra and an increase in inter population genetic diversity of *P. splendens* in the future, and therefore a potentially increase of vulnerability to extinction of remaining populations.

### Conservation status

Our assessment of the level of threat of *P. splendens* using the IUCN criteria revealed that based on our data, this species should be reclassified as Endangered. Our results differ from Hechenleitner et al. (2005) who classified *P. splendens* as critically endangered (B1ab(iii)). Unfortunately, we cannot compare our results to Hechenleitner et al. (2005) as they did not report what information they used to perform their assessment.

We could not calculate or find reliable data to estimate the area of occupancy of the species. Although there is data from the national forest inventory that could help to estimate an area of occupancy, it seems that the inventory did not include accurate data at the species level, and only three sites containing *P. splendens* were apparently mapped, representing an extension no larger than 400 ha. In contrast, based in our field survey we presume that the area of occupancy is not smaller than 2.000 ha. However, even taking an optimist approach of 3.000 ha for the area of occupancy, this would not change *P. splendens* to any other conservation status (e.g. from endangered to vulnerable) because other assessment criteria will prevail.

We had a similar challenge for estimating the size of the remaining populations. We only could calculate densities of *P. splendens* based in roughly estimation performed in each of the visited sites. Thus, these estimations corresponded to projected abundance for the specific visited site, but do not reflect an estimation for the entire population at that point, and therefore they cannot be used to calculate the total number of individuals present in the range of distribution. However, as happened with the area of occupancy variable, the classification of *P. splendens* as endangered is not based in the lower number of individuals we were able to estimate, but in other assessment criteria. Therefore, even though further studies are able to estimate larger population’s size, the threat level of *P. splendens* would not change.

## Conclusion

Results from our work provide new and essential information for assessing current conservation status of *P. splendens* populations. The integration of our results (i.e., niche modelling, land-use change estimation, and genetic variability analysis) suggest that *P. splendens* had a large continuous population that extended through the entire range of historical recorded populations, providing essential evidence to support the continuous population assumption made previously by other authors (e.g., Hechenleitner et al. 2005; Muñoz & Serra 2006).

We were also able to update information regarding the current populations, establishing that there are only two main areas with *P. splendens* populations left, as was also suggested by previous studies. However, we found that these remaining populations are more reduced in extension and density that was previously thought. Moreover, the larger and better conserved of these populations (i.e., Pichidangui-Los Molles) is currently heavily threatened by anthropic pressure, including goat grazing, exotic tree plantations, illegal trash dumping, increasing fires, and real-estate developments. Although this area has been declared as an Urgent Priority Site for Conservation by the Chilean government (CONAMA-PNUD 2005), until now, there has not been any official conservation action taken. More concerning is the fact that currently the local government is soon to officialize a new urban development normative, which is proposing to transform more than 1700 ha of the Los Molles-Pichidangui *P. Splendens* population habitat, from natural lands, to residential areas (Seremi Medio Ambiente Valparaíso 2014). Because this probable scenario, *P. splendens* may soon change from endangered to critically endangered.

Based on the results from our work we recommend: (1) update the information available to the IUCN and evaluate the reclassification of *P. splendens* as Endangered, (2) take urgent public and private actions to conserve the remaining populations of *P. splendens*; especially the larger populations which is now threatened by the upcoming change in the land use normative, and (3) develop more studies to fill the knowledge gap regarding this species.

## Competing interests

The authors declare that they have no competing interests.

## Authors’ contributions

NM and IF conceived the study objectives, design and methodologies, and were in charge to draft the manuscript and coordinate inputs from the other co-authors. NM performed the niche modelling analysis. IF performed the land-use change analysis and mapping. BC carried out the genetic analysis and helped with drafting later version of the manuscript. CO helped with the literature review and in drafting the manuscript. All the above authors have read the manuscript and approved this final version.

## Acknowledgments

We would like to thank Miguel Gómez from the Faculty of Agronomy and Forestry Engineering of the Pontifical Catholic University of Chile for his assistance in recollecting historical presence data. We also would like to thank Giselle Muschett and Michelle Jones for their help polishing the last version of the manuscript. Fieldwork and genetic analysis were funded by the Chilean Native Forest Research Grant, Project CONAF 025/2010: “Distribución, hábitat potencial y diversidad genética de poblaciones de Belloto del Norte (*Beilschmiedia miersii*) y Lúcumo Chileno (*Pouteria splendens*)”. Plant material extraction and sampling methods were authorized by the Chilean National Forest Corporation (CONAF).

## Supplementary material

**Supplementary material 1.** *Pouteria splendens* population presence records information. Sites are listed from north-to-south. Sites in bold type represent those for which we were able to find individuals. Approximate numbers of individuals are only presented for populations found during fieldwork. Numbers are based in visual estimations from fieldwork and recent satellite images. ^a^This site was not in the historical records and was registered during the field work. ^b^This was considered as an spatial outlier point and was not used in the modeling process because corresponded to a registry from 1926 for which we did not have enough reliance.

| Site name | Coordinates |  | Approx. number of individuals | Herbarium |
| --- | --- | --- | --- | --- |
|  | Latitude | Longitude |  |  |
| Quebrada el Buitre | 31°26'52"S | 71°35'3"W |  | CONC |
| Norte de Los Vilos | 31°41'16"S | 71°31'45"W |  | CONC |
| La Totora | 31°54'30"S | 71°27'41"W |  | CONC |
| Estación Ingeniero Barriga | 32°1'57"S | 71°29'54"W |  | CONC |
| Sur de Los Vilos | 32°3'20"S | 71°30'54"W |  | CONC |
| Entre Pichidangui-Los Vilos | 32°3'53"S | 71°31'4"W |  | CONC |
| **Fundo Palo Colorado** | 32°4'17"S | 71°29'24"W | 60 | CONC |
| **Fundo Palo Colorado Oeste<sup>a</sup>** | 32°5'34"S | 71°30'55"W | 50 | <sup>a</sup> |
| Silla del Gobernador | 32°7'50"S | 71°28'52"W |  | SGO |
| **Pichidangui** | 32°8'53"S | 71°31'38"W | 300 | CONC |
| **Los Ermitaños** | 32°11'0"S | 71°31'46"W | 500 | CONC |
| **Kilómetro 180 ruta 5 norte** | 32°12'35"S | 71°30'41"W | 200 | CONC |
| **Los Molles** | 32°14'9"S | 71°31'18"W | 400 | CONC |
| **Quebrada el Chivato** | 32°13'59"S | 71°29'33"W | 50 | CONC |
| Quebrada Las Docas | 33°2'16"S | 71°39'17"W |  | CONC |
| Las Lúcumas | 33°2'22"S | 71°38'56"W |  | CONC |
| Valle del Agua Potable | 33°2'33"S | 71°38'42"W |  | CONC |
| La Palma<sup>b</sup> | 32°58'32"S | 71°16'31"W |  | SGO |
| Curauma-Las Docas | 33°4'27"S | 71°39'35"W |  | CONC |
| Laguna Verde | 33°6'15"S | 71°40'17"W |  | CONC |
| **Curaumilla** | 33°8'8"S | 71°42'21"W | 20 | CONC |
| **Caleta Quintay** | 33°12'18"S | 71°41'53"W | 70 | SGO |
| Algarrobo | 33°22'53"S | 71°40'51"W |  | CONC |

**Supplementary material 2.** Corrected Akaike information criterion (AICc) for the best 10 combinations of features and regularization multiplier values of MaxEnt (e.g LQ 1 = linear quadratic and regularization multiplier equal to 1).* Auto features corresponds to using MaxEnt with the default configuration parameters for small samples.

| Combination of features | AICc score |
| --- | --- |
| LQ 1 | 421.17 |
| L 1 | 428.23 |
| LQ 2 | 429.08 |
| L 2 | 441.58 |
| LQ 5 | 447.04 |
| LQH 5 | 455.14 |
| Auto features* | 457.90 |
| LQH 1 | 457.90 |
| LQH 2 | 465.54 |
| H 5 | 468.05 |

**Supplementary material 3.** ISSR primers used for the analysis of *Pouteria splendens*. R can be either A or G nucleotide, whereas Y can be either C or T nucleotide

| Primer | Sequence | Annealing temp (C) | Band size range (base pairs) |
| --- | --- | --- | --- |
| 825 | ACA CAC ACA CAC ACA CT | 58 | 500-2700 |
| 844 | CTC TCT CTC TCT CTC TRC | 54 | 420-1900 |
| 849 | GTG TGT GTG TGT GTG TYA | 57 | 410-1700 |
| 854 | TCT CTC TCT CTC TCT CRG | 54 | 430-2080 |
| 856 | ACA CAC ACA CAC ACA CYA | 59 | 430-2080 |

**Supplementary material 4.**
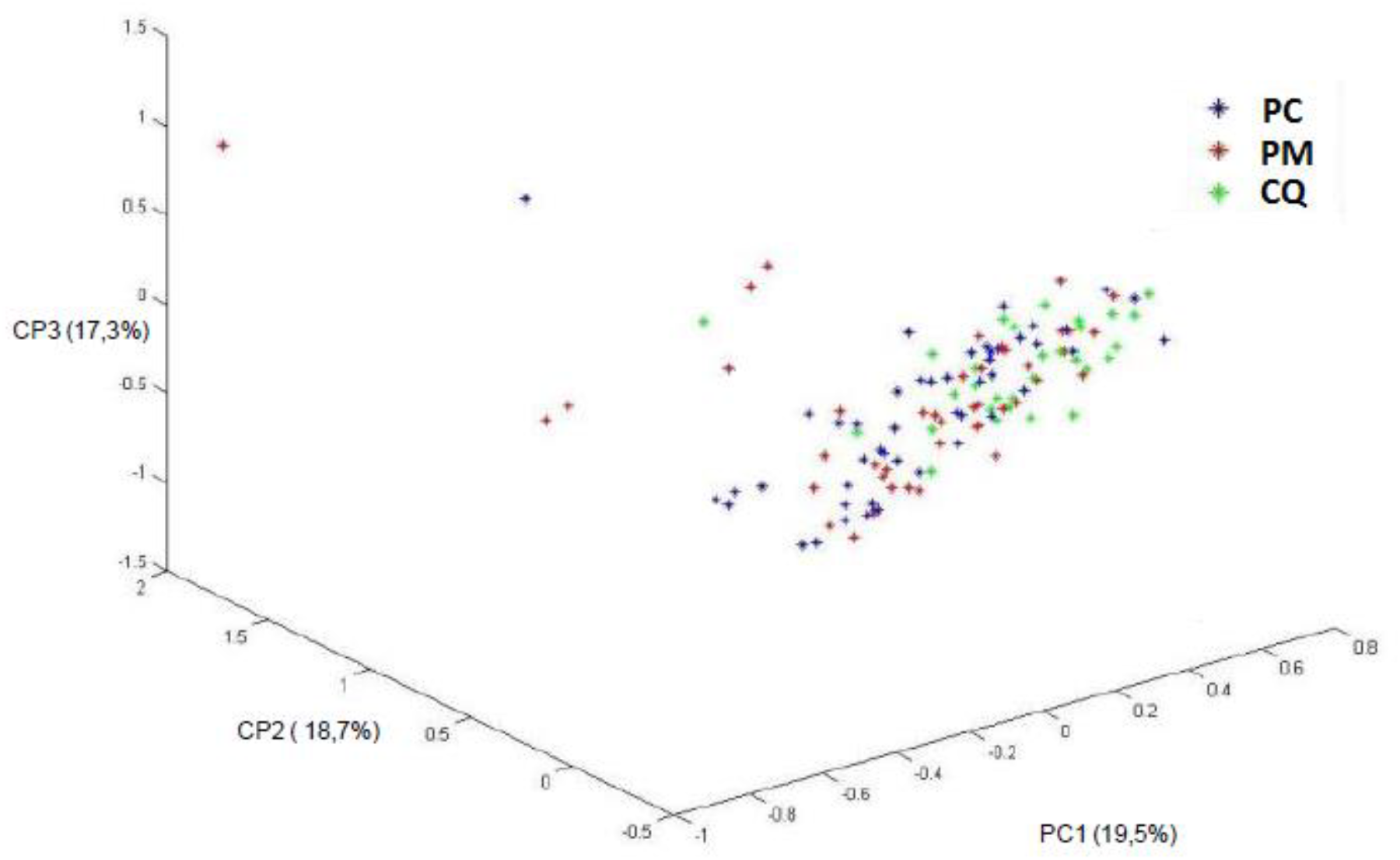
Principal Component Analysis for 121 individuals of *P. splendens* and 147 ISSR loci. Sampled populations were PC (Palo Colorado), PM (Pichidangui-Los Molles), CQ (Caleta Quintay).

